# Coral settlement module designs for scalable reef restoration

**DOI:** 10.64898/2026.06.24.733534

**Authors:** J Reichert, M Asbury, R Argall, GK Chen, J Ehrenberg, Z Huang, B Jones, H Jorissen, J Levy, AD Nims, ME Rottmueller, LH Rova, A Thode, D Wangpraseurt, the R3D Consortium, JS Madin

## Abstract

The global coral reef crisis has prompted restoration initiatives worldwide. Targeting the coral larval stage is among the most scalable approaches as recruitment operates over large spatial scales. It thus represents one of the best levers for coral population recovery. Active coral larval seeding has shown considerable success, and passive substrate engineering has emerged as a promising complementary strategy. Coral settlement modules featuring helix recesses have increased settlement and survival by up to 80-fold on small experimental units, but whether these results translate to tools deployable at the scale of thousands of units, remains yet an open question. Here, we transferred structural features from successful experimental coral settlement designs into production-ready concrete modules to (i) evaluate coral recruitment on five designs at four reef sites differing in flow regime and coral cover over one year; (ii) compare production-scale performance against experimental clay modules and natural reef substrate; and (iii) identify key parameters for large-scale production. The helix recess geometry of coral settlement modules outperformed the featureless control design approximately 20-fold and exceeded natural reef recruitment at least 3- to 32-fold. The helix features were successfully transferred from experimental clay to production-scale concrete modules, yielding comparable settlement densities when standardized to crevice length, which proved to be the biologically relevant unit of available habitat. Production feasibility was demonstrated by producing 690 modules for deployment on a hybrid reef on the west side of Oʻahu, Hawaiʻi. The passive coral larval recruitment approach presented here could substantially improve the logistical and economic feasibility of large-scale coral reef restoration. This approach requires neither coral larval rearing, handling, nor coral fragmenting, and is compatible with active larval seeding where genetic diversity or larvae supply are limiting factors. The coral settlement modules can be cast in standardized concrete molds at precast facilities. Modules have demonstrated consistent coral recruitment enhancement across reef environments with contrasting flow and coral cover. Deploying mixed arrays of helix-recess structures with designs offering multi-level complexity and three-dimensional rugosity maximizes outcomes for coral, fish, and invertebrate communities simultaneously. Site selection is the most critical deployment decision and should consider larval supply, hydrodynamics, and substrate stability which drive recruitment outcomes more than design choice alone. The modules offer a range of application potential, ranging from integration into existing coastal infrastructure over stand-alone reef restoration approaches, to substrate-consolidating interconnected arrangements.

## Introduction

The global coral crisis has led to the loss of 30–50% of reef cover over recent decades, driven by climate change and local anthropogenic stressors (Carpenter et al. 2008; Hughes et al. 2017; IPCC 2023). This trend has prompted restoration initiatives worldwide, aiming to combat habitat loss by restoring structural complexity or boosting reef community resilience (Boström-Einarsson et al. 2020; Randall et al. 2020; Suggett & van Oppen 2022). Many approaches focus on coral transplantation. However, the outplanting of fragments is labor- and cost-intensive and requires the sacrifice of donor colonies (Boström-Einarsson et al. 2020). Restoration approaches have therefore increasingly targeted early life-history stages of corals (Boström-Einarsson et al. 2020). A single coral colony can release millions of larvae during a spawning event (Harrison & Wallace 1990; Hall & Hughes 1996), making coral larval recruitment one of the most scalable levers for restoration (Madin et al. 2025).

Active larval seeding has successfully enhanced coral recovery on degraded reefs. For example, the delivery of *Acropora tenuis* larvae in mesh enclosures achieved 2.3–5.7 m⁻² colonies after three years in the Philippines (Cruz & Harrison 2017; Harrison et al. 2021). The CSIRO Larval Seedbox enhanced settlement up to 56-fold compared to background rates across 2 ha at Lizard Island (Doropoulos et al. 2025). Settlement structures seeded with larvae and outplanted onto the reef have retained at least one live coral in 24–60% of devices across hectare-scale Great Barrier Reef deployments (Chamberland et al. 2017; Ramsby et al. 2026). However, their success strongly depends on habitat properties. Sand deposition and unconsolidated substrate can reduce recruit survival to near-zero despite high initial settlement (Waters et al. 2025; Ramsby et al. 2026), making site assessment critical prior to deployment.

Furthermore, passive approaches have been developed to enhance larval settlement *in situ*. For example, rubble stabilization creates consolidated surfaces for wild larval settlement (Ceccarelli et al. 2020; Leung et al. 2025). Other passive approaches are informed by the underlying mechanisms that identify surface topography, substrate material, hydrodynamics, and biological cues strongly influence larval settlement and survival (Whalan et al. 2015; Hata et al. 2017; Jorissen et al. 2021; Levenstein, Gysbers, et al. 2022; Levenstein, Marhaver, et al. 2022; Edmunds 2023). Within these parameters, mesoscale structural design has emerged as a key determinant of both settlement and post-settlement survival (Doropoulos et al. 2016; Randall et al. 2020). This is because coral recruits experience extremely high early mortality, and micro-crevice structures provide physical refuge from predation and grazing (Doropoulos et al. 2016; Nozawa 2008), while larvae actively select settlement positions based on crevice geometry, depth, and light exposure (Hata et al. 2017). Building upon this, engineered settlement environments have been explored as a passive restoration approach.

An investigation of crevice design principles showed that angled spiral crevices outperform vertical crevices for both coral settlement and post-settlement survival, and accumulate greater benthic richness driven by cryptic diversity (Schiettekatte et al. in review). Preferred recess properties were identified in a follow-up field study, testing seven 3D-printed ceramic modules incorporating different structural features, such as helix recesses of varying depths, widths, and angles (Reichert et al. 2025). Structures with helix recesses dramatically outperformed all other designs, increasing settlement approximately 80-fold and post-settlement survival 20- to 50-fold over one year compared to controls, and approximately 70-fold compared to natural reef substrates (Reichert et al. 2025). Subsequent work demonstrated functional robustness across geometries and materials when helix recesses were transferred to cylindrical concrete structures (Reichert et al. 2026). Together, these studies establish helix recesses as a generalizable microhabitat principle with the potential for engineering applications at reef scale.

Despite the demonstrated effectiveness of helix recesses at the experimental scale, a critical gap remains between proof-of-concept and scalable deployment. Structures tested to date have been small (10–20 cm diameter), individually produced by 3D clay printing, and deployed under research conditions. Scaling to the reef level requires larger structures that can be manufactured rapidly in volume from durable, reef-compatible materials suited for industrial fabrication (Boström-Einarsson et al. 2020; Higgins et al. 2022; Ramsby et al. 2026). Key unknowns include the fidelity of mold-based casting for complex geometries and biological performance across reef habitats differing in water flow conditions and coral cover.

Therefore, the goal of this study was to transfer structural features identified as successful in experimental coral settlement structures to production-ready, upscaled designs, and to evaluate their performance across coral reef habitats. Specifically, we (i) evaluated recruitment on five structural designs at four reef sites differing in flow regime and coral cover over one year; (ii) compared recruitment on production-scale modules with experimental helix recess designs and adjacent natural reef substrate; and (iii) identified key parameters for large-scale module production.

## Methods

### Experimental design

To test whether the transfer of structural features promoting coral settlement and survival to production-ready modules was successful, we evaluated the performance of four large format concrete coral settlement module designs (Fig. 1; Control Dome, Hexagon Dome, Fractal Dome, and Superdome; diameter of 50 cm, height of 25 cm). These were tested in a field experiment at four locations on the windward side of Oʻahu, Hawaiʻi (Patch Reef 13: 21.4504°N, 157.7956°W; Hinalea Reef: 21.4324°N, 157.7869°W; Lighthouse Reef: 21.4322°N, 157.7900°W; and Ulupaʻu Reef: 21.4444°N, 157.7352°W) which differ in coral cover and flow regime. Coral cover at each site was estimated from Google Earth satellite imagery using unsupervised image classification with the ISO Cluster algorithm in ArcGIS Pro 3.6.1. The imagery was first classified into 15 classes, after which reef-associated classes were manually selected to calculate percent cover (Figure S1). Sites were categorized as high (≥60%, Patch Reef 13 and Barrier Reef), medium (20–60%, Hinalea Reef and Lighthouse Reef), or low (<20%, Ulupaʻu Reef) coral cover. Flow regime categories were assigned during site visits for deployment and monitoring, in relation to known flow regimes within the area (Lowe et al. 2009) as high (Patch Reef 13, Ulupaʻu Reef, and Barrier Reef), medium (Hinalea Reef), or low (Lighthouse Reef) flow. The Superdome was deployed at all four sites to assess performance under varying habitat conditions. The full design comparison among all four module types was conducted at two sites (Lighthouse Reef and Ulupaʻu Reef). Additionally, the large-format concrete Superdome was compared to small experimental-scale clay and concrete Superdomes at Patch Reef 13 to evaluate the effect of scale and material. Coral recruitment and survival were monitored at 2 months, 6 months, and 1 year post-spawning at each site. Recruitment on settlement modules was compared to natural reef recruitment at the Barrier Reef in Kāneʻohe Bay at the 1-year time point, which was chosen as the natural recruitment comparison. The Barrier Reef is adjacent to Patch Reef 13, receives high larval supply from nearby fully covered reefs, and offers hard substrate not saturated by live coral cover, making it a suitable benchmark for maximum recruitment in a high-quality reef environment within the bay. Field activities involving coral reef organisms at Kāneʻohe Bay were conducted under the Hawaiʻi Institute of Marine Biology Special Activity Permit 2024-45. Activities at Ulupaʻu, Mōkapu Peninsula were authorized under DLNR Office of Conservation and Coastal Lands Site Plan Approval OA 24-08, NOAA NMFS Endangered Species Act Section 7 and Essential Fish Habitat consultations (I-PI23-2203-DG; PIRO-2023-02584), and a NEPA Categorical Exclusion (12030203.A.17 and 12030203.A.19).

**Figure 1:**
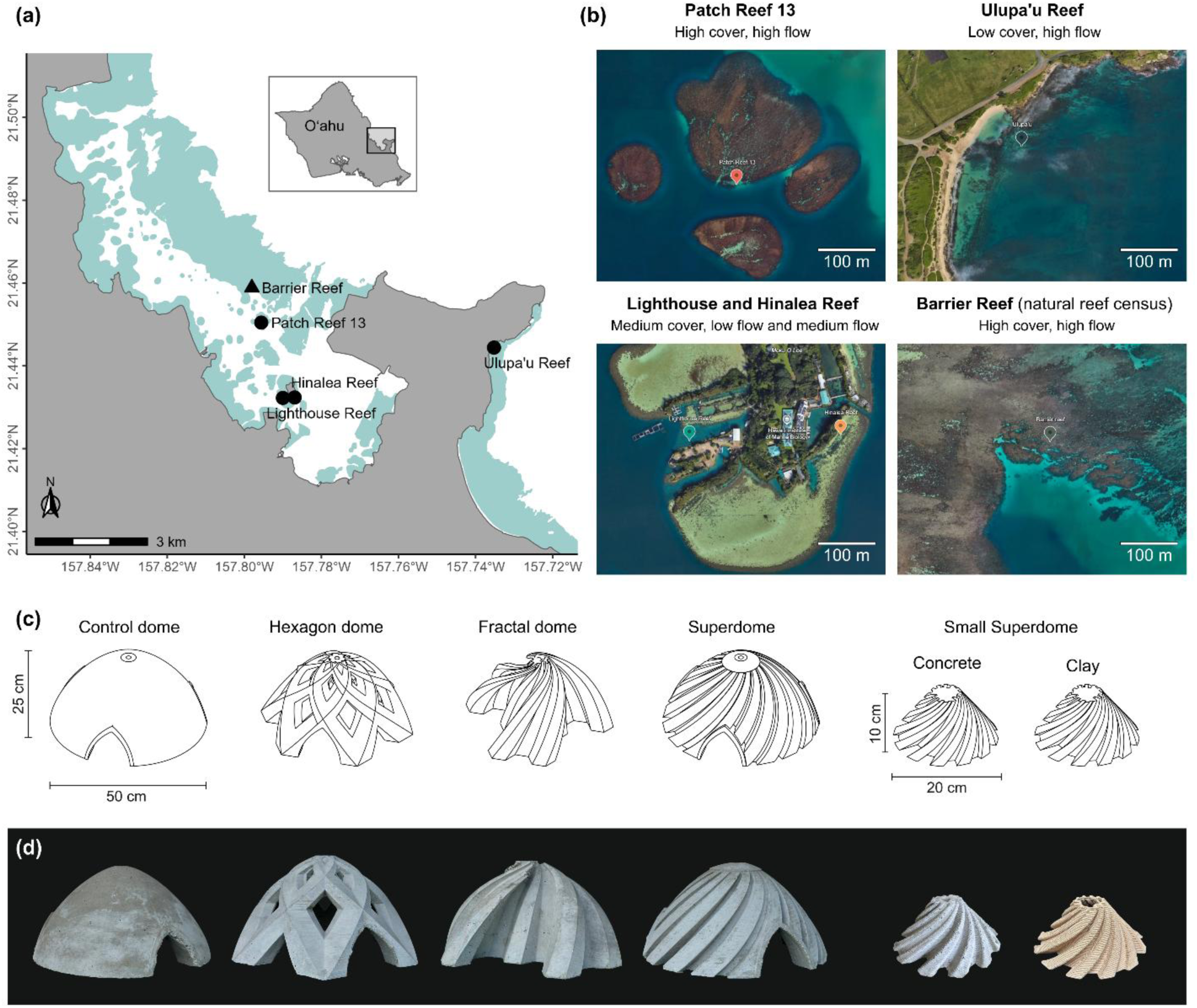
Study locations and coral settlement module designs. (a) Map of study locations on the windward side of Oʻahu, Hawaiʻi. Modules were deployed at four locations (circles, Patch Reef 13, Hinalea Reef, Lighthouse Reef, Ulupaʻu Reef) and natural recruitment was assessed as a comparison at the Barrier Reef (triangle). Light green areas indicate the presence of reef. (b) Satellite images of the four deployment locations. Labels indicate the flow regime and coral cover category of each site. (c) Technical drawings of the five coral settlement module designs: four large concrete modules (Control dome, Hexagon dome, Fractal dome, and Superdome; 50 cm diameter, 25 cm height) and the Small superdome (20 cm diameter, 10 cm height), produced in both concrete and clay as a size- and material-matched comparison with the successful experimental helix recess design. (d) Photographs of the corresponding coral settlement modules.

### Coral settlement module design and production

Coral settlement modules were designed in Blender (v3.4), modifying previous designs from Reichert et al. (2025). Models were further processed in Autodesk Fusion (v27) to integrate an internal cave system and design mold components for production. The design rationale for the upscaling was to keep the recesses at the same absolute dimension as the proven experimental design rather than scaling them up with the larger module leading to wider and deeper crevices, because coral recruits respond to the absolute, millimeter-scale geometry of the recesses rather than to overall module size. The effective recess depths and widths identified in the experimental modules were therefore retained, and only the shallowest recess category, which performed worst in earlier work, was omitted. The smaller recesses were maintained at the tops of the domes, where they are least likely to be affected by sediment accumulation when deployed on the seafloor. In addition, the original conical design was modified to a rounded dome shape to minimize potential interference with other marine organisms by reducing sharp edges and creating a smoother external profile.

All large modules were 50 cm in diameter and 25 cm in height. Four designs were created (Fig. 1): (1) The ’Superdome’ incorporated 18 helix recesses of three different depths (20–40 mm at the base, tapering to 2.5–6 mm at the top), and widths of 30 mm at the base tapering to 7 mm at the top; (2) the Hexagon Dome which provided openings of three size classes; (3) the Fractal Dome, which provided exposed, sheltered, and cryptic microhabitats through deeply recessed surfaces; and (4) the Control Dome, a smooth dome without surface features. The shallowest recess category previously tested in the experimental-scale Small Superdome was excluded from the production-scale Superdome because it consistently yielded the lowest recruit densities in a previous study (Reichert et al. 2025). Detailed dimensions of all module features are provided in Figure S2 and Table S1.

Upscaled modules were produced via concrete casting (Fig. 2). Module positives were 3D printed from PLA plastic (BigRep Bio/Classic, 2.85 mm diameter) using a large-format fused deposition modelling (FDM) printer (Big Rep ONE, Big Rep GmbH, Berlin, Germany) equipped with a PEX Fiber Hotend (1 mm nozzle diameter, 0.6 mm layer height). Module positives were printed with a wall thickness of 3 mm, 15% grid infill, and buildplate support structures, at a nozzle temperature of 200°C and a bed temperature of 60°C. For each settlement module design, a flexible silicone mold was created by pouring Mold Star 30 platinum-cure silicone rubber (Smooth-On, Inc., Macungie, Pennsylvania, USA) into a 3D-printed outer shell holding the module positive and a cave inset, and kept in place by a 7/16-inch threaded rod. After curing overnight, the mold was disassembled, and the module positive out of PLA plastic was removed. With the mold now being ready to cast concrete domes, the mold was reassembled around the cave inset, and modules were cast using a high-strength concrete mix (Quikrete Concrete Mix No. 1101, Quikrete Companies, Atlanta, Georgia, USA). Modules cured in the mold for two days before removal. No pH buffering or pre-soaking treatment was applied before deployment, and any surface alkalinity effects are assumed to be consistent across all module types. Modules were deployed one to two weeks before the predicted spawning event, providing additional time for surface pH to equilibrate with ambient seawater prior to larval settlement. Finished modules weighed between 14 and 27 kg, depending on design. All tested modules remained structurally stable following the curing protocol and deployment over the course of one year.

**Figure 2:**
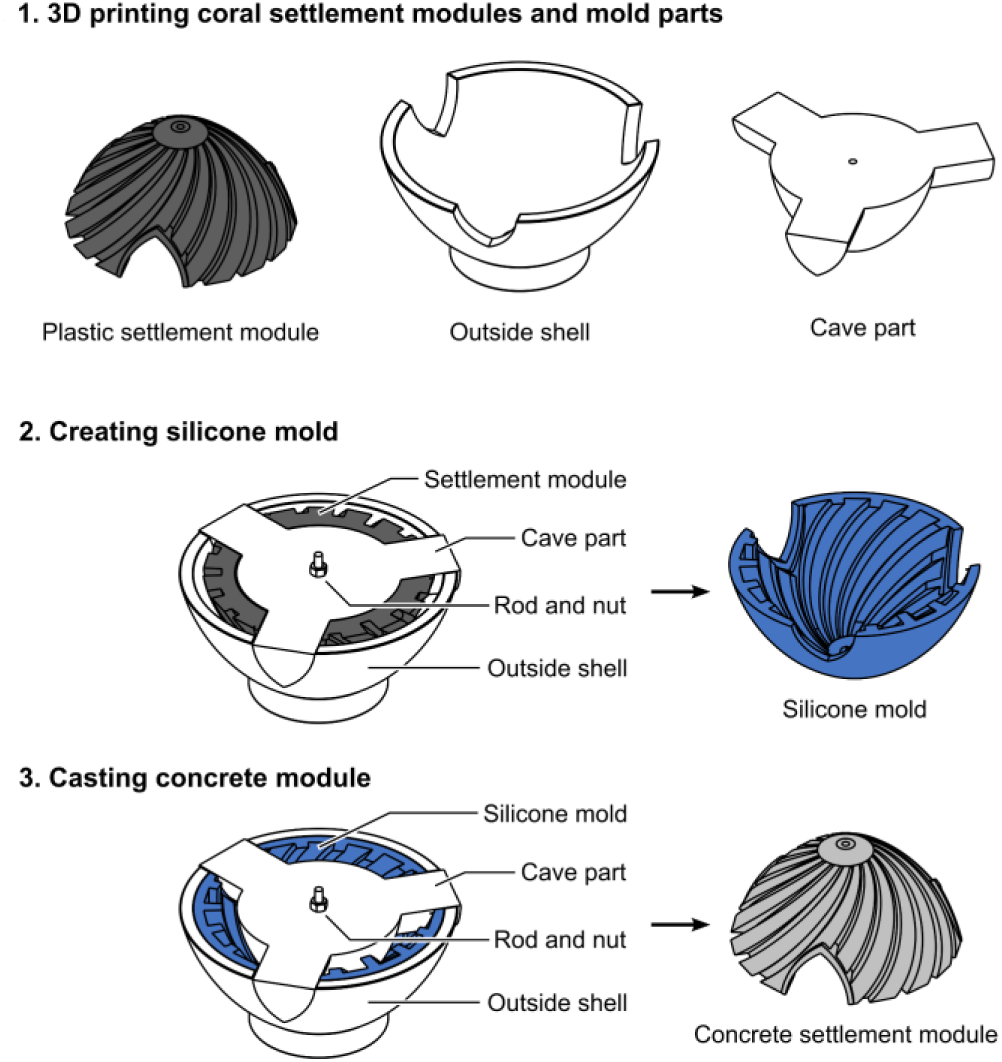
Production steps of casting large concrete coral settlement modules. Module designs and mold parts (outside shell and cave part) were 3D printed in plastic (PLA). The module positives and mold parts were assembled and connected via a threaded rod and hex nuts (7/16“). A silicone mold was created for every settlement module design. The concrete domes were cast using the created silicone molds combined with the 3D printed mold parts.

The experimental-scale Small Superdome modules (20 cm diameter, 10 cm height) held 12 helix recesses spanning four depth categories (10–40 mm at the base, tapering to 2–11 mm at the top). Crevice widths tapered from 30 mm at the base to 6 mm at the top. Small Superdomes were produced in two materials. Clay modules were 3D printed with a Delta Wasp 40100 3D printer (WASP, Italy) using a mid-fire pottery clay (Soldate 60, Aardvark Clay and Supplies, USA) extruded through a 4 mm nozzle, producing a layered surface microstructure. Printed modules were air-dried in a stable, air-conditioned environment for 5–10 days until bone-dry, then fired in a kiln (Jupiter Sectional Kiln, L&L Kilns, USA) at approximately 1100°C (cone 8). Concrete Small Superdomes were cast using a silicone mold made from a fired clay dome and the same concrete mix was used as for the large modules (Quikrete Concrete Mix No. 1101). The silicone mold preserved the original dimensions and surface microstructure of the clay module, so both materials shared the same geometry.

Five modules included in the experiment carried minor design modifications. Three Superdomes at Hinalea Reef had 0.5 cm holes drilled at 10 cm intervals along the base of the recesses to accommodate an Underwater Zooplankton Enhancement Light Array (UZELA™) underwater light, which illuminated one-third of the dome during the first 2–3 months of the experiment (Jorissen et al. in review). Since the light treatment did not show a significant effect on settlement or survival, the structures were included here for comparison. Two Superdomes at Lighthouse Reef had plug attachment holes on the outer recess surface, for testing the ability to add coral fragments to the modules (no fragments were added to the studied modules). These areas were excluded from the recruit counts to avoid introducing additional structural complexity and allow for comparison with the other modules.

### Module deployment and monitoring of coral recruitment

Coral settlement modules were deployed at four sites on the windward side of Oʻahu one to two weeks before the predicted *Montipora capitata* spawning on July 5, 2024 (Fig. 3a). At Patch Reef 13 and Hinalea Reef, modules were mounted on experimental tables. At Lighthouse Reef and Ulupaʻu Reef, modules were placed directly on the seafloor. Both on tables and on the seafloor, the weight of the modules (14.0–27.1 kg) was generally sufficient to keep them in place without additional anchoring. The exception was Ulupaʻu Reef, where an unusually strong swell displaced a few modules, indicating that anchoring may be required at more energetic sites.

**Figure 3:**
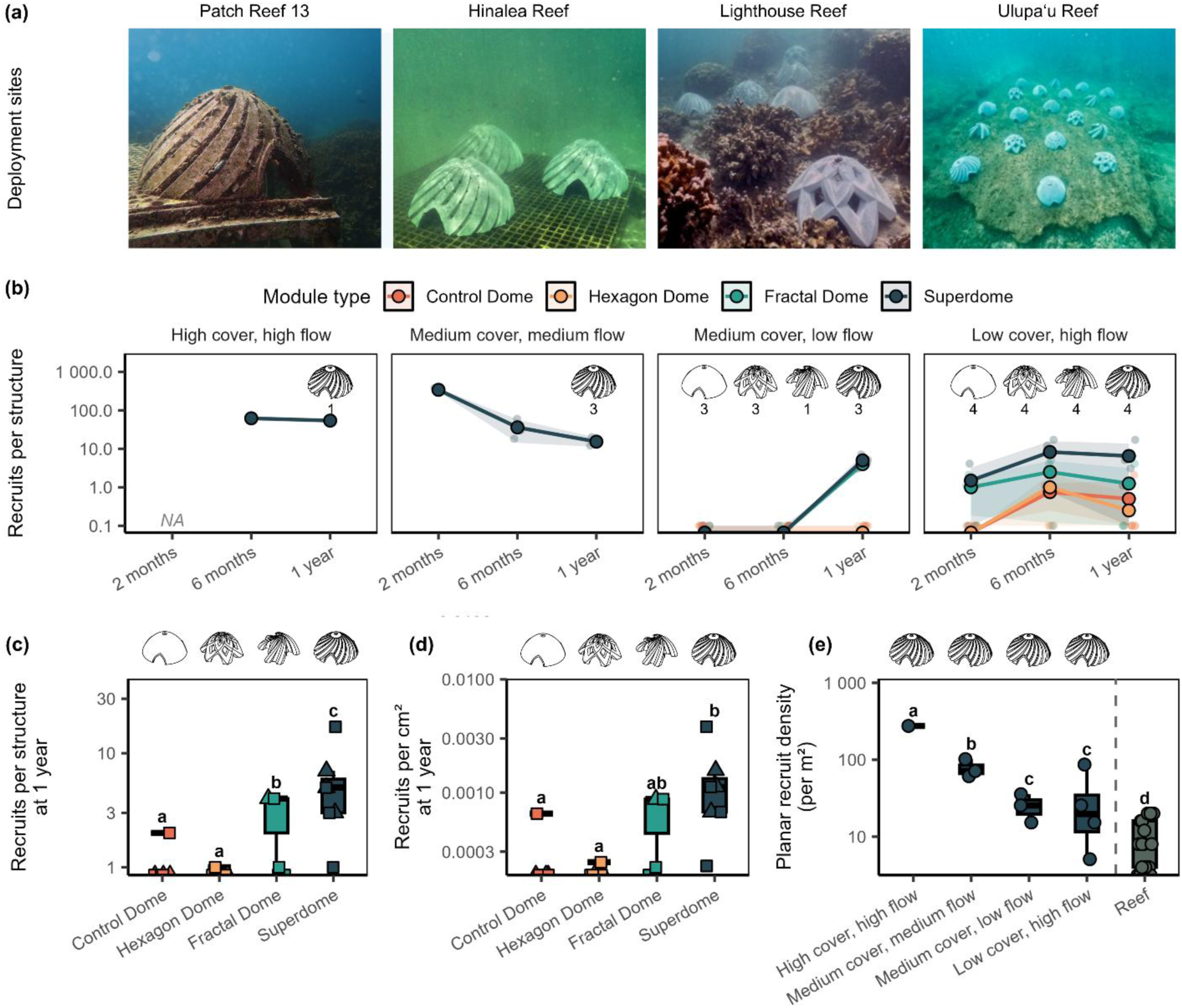
Coral recruitment on coral settlement modules across four reef environments. (a) Overview photo of deployed domes *in situ* at the four test locations (Patch Reef 13 after 1 year, and Hinalea Reef, Lighthouse Reef, and Ulupaʻu Reef at deployment). (b) Recruits per structure for four module designs (Control, Hexagon, Fractal, and Superdome) across all four study sites (differing in coral cover and flow regime) at 2 months, 6 months, and 1 year post-spawning. (c,d) Recruits per structure and recruits per cm² of module surface area for the two fully replicated sites, Lighthouse Reef (triangles) and Ulupaʻu Reef (squares), at 1 year. Letters denote significant differences from BH-adjusted pairwise emmeans contrasts of a negative-binomial GLMM with site as a random effect. (e) Coral recruitment on Superdome modules across the four reef environments with contrasting coral cover and flow (Patch Reef 13, Hinalea Reef, Lighthouse Reef, Ulupaʻu Reef, from left to right), compared to reef counts on natural reef substrate in the Barrier Reef in Kāneʻohe Bay. Recruitment is compared as planar recruit densities. Letters denote significant differences from BH-adjusted pairwise emmeans contrasts of a negative-binomial GLM. Note: Hinalea Reef data (medium cover, medium flow) were collected *ex situ* under white light on dried surfaces at 1 year and may overestimate recruit density relative to other sites.

The number of modules *n* deployed per design and site varied due to logistical constraints in production prior to the spawning timing (Table S2). The Superdome was deployed at all four sites (Patch Reef 13: n = 1; Hinalea Reef: n = 3; Lighthouse Reef: n = 3; Ulupaʻu Reef: n = 4). The full four-design comparison was conducted at Ulupaʻu Reef (n = 4 per design) and Lighthouse Reef (Control Dome: n = 3; Hexagon Dome: n = 3; Fractal Dome: n = 1; Superdome: n = 3). The size and material comparison was conducted at Patch Reef 13, where the large concrete Superdome (n = 1) was compared with small concrete Superdomes (n = 3) and small clay Superdomes (n = 4). Hinalea Reef contained only Superdomes (n = 3), which was part of another experiment (see Coral settlement module design and production) and therefore contributed to the across-site Superdome comparison but not to the four-design comparison.

Coral recruitment on modules was assessed at 2 months, 6 months, and 1 year post-spawning. Recruits were detected *in situ* while snorkeling using blue-light fluorescence (Piniak et al. 2005; Sola Nightsea blue light with blue-light-blocking glasses) and counted separately on exposed outer surfaces and within recesses. An exception was made for the Hinalea Reef Superdomes at the 1-year census, which were retrieved and counted *ex situ* under white light on dried surfaces due to logistical constraints of conducting the in-water survey. White light census of dried surfaces may overestimate recruit counts relative to *in situ* blue-light censuses, as non-fluorescing or dead corallites may also be recorded. Counts from this time point should therefore be compared with caution to *in situ* blue-light census data from other sites.

Recruit counts were converted to densities (recruits per cm²) by standardizing to the outer surface area of each module type. For the Superdome modules (large and small), recruit numbers were additionally standardized to planar area (recruits per cm²), recess surface area (recruits per cm²), and recess length (recruits per cm) to facilitate comparison across locations and with smaller experimental units from previous studies.

Natural reef recruitment was assessed at the Barrier Reef in Kāneʻohe Bay (21.4589°N, 157.7980°W; high coral cover, high flow) one year after spawning. Coral recruits were counted by SCUBA divers in 18 randomly selected 0.5 × 0.5 m quadrats using blue light fluorescence. Quadrat locations were identified within a three-dimensional photogrammetric reconstruction of the site following established procedures (Pizarro et al. 2017), processed in Agisoft Metashape (Roach et al. 2021), and their positions were printed on paper for use during subsequent in-water surveys. Bootstrap subsampling analysis confirmed that the mean recruit density estimate stabilized well before n = 18 quadrats, indicating adequate sample size for characterizing the site-level baseline. Recruitment density was standardized to the planar surface area of each quadrat (0.25 m^2^).

### Large-scale production

The upscaled module designs were mass-produced for a large-scale field deployment as part of the Rapid Resilient Reefs for Coastal Defense (R3D) program, a three-year, $27 million initiative funded by the Defense Advanced Research Projects Agency (DARPA) through the University of Hawaiʻi. The project aimed to develop self-healing hybrid biological and engineered reef structures for coastal protection and wave attenuation (DARPA Reefense program; DARPA 2024). Meeting the production demands of R3D required the transfer of the experimental module designs to a precast concrete manufacturing workflow capable of serial production of modules at volume. Master patterns for each module design were 3D printed from ZYLtech PETG filament using Modix Big Meter or Big 60 printers equipped with a Supervolcano hotend (1 mm nozzle diameter, 0.6 mm layer height), sharing the same CAD geometry as the original experimental mold positives. Production molds were created from these master patterns, and modules were cast at Jensen Infrastructure (Kapolei, Oʻahu, Hawaiʻi; jensenprecast.com) at a rate of one dome per mold per day.

### Statistical analysis

All statistical analyses were performed in R (v4.5.0; R Core Team 2025). Since the recruit count data were overdispersed, negative binomial (NB) generalized linear mixed models (GLMMs) were used to analyze count data, fit via the glmmTMB package (Brooks et al. 2017). Model terms were evaluated using Type II Wald chi-square tests. Pairwise contrasts among factor levels were computed using the emmeans package (Lenth et al. 2025), and p-values were adjusted for multiple comparisons using the Benjamini-Hochberg (BH) false discovery rate procedure throughout.

Recruit numbers and densities on the different settlement module designs were compared at Lighthouse and Ulupaʻu Reef after 1 year, the only two sites where all four dome types were deployed. A negative binomial GLMM (NB2 parameterization; glmmTMB) with dome type as a fixed effect and dome identity nested within location as a random intercept was fitted. NB2 was preferred over NB1 for its better accommodation of strongly overdispersed counts. Differences among dome types were evaluated using a Type II Wald chi-square test, and BH-adjusted emmeans pairwise contrasts identified differences among dome types. To test whether recruitment densities differed, the same model included the log of module surface area as an offset, standardizing estimated marginal means to recruits per cm².

To compare the performance of coral settlement modules across reef environments and against natural reef substrate, planar recruit densities were compared. For this, we combined Superdome recruit counts from all four sites with natural reef counts from the Barrier Reef. A negative binomial GLM was fitted with reef environment as a fixed effect and log planar area as an offset, accounting for differences in sampled area across substrates and module types.

To assess large concrete Superdome performance relative to small clay and concrete Superdomes at Patch Reef 13, a hurdle approach accounted for crevices containing no recruits. Crevice occupancy (binary, any recruits present or not) was modeled with a binomial GLMM (logit link), and recruit density among occupied crevices with separate Gamma GLMMs (log link) for recess-area density (recruits cm⁻²) and perimeter density (recruits cm⁻¹). In all models, crevice depth was a fixed covariate and dome identity a random intercept. The small number of domes per type (n = 1–4) limits reliability of random effect estimates; results should be interpreted accordingly. All hurdle components were fit via lme4 (Bates et al. 2015), and the planar negative binomial GLM was fit via glmmTMB. Crevices on outside dome surfaces were excluded. The effect of crevice depth was additionally evaluated in a Gamma GLMM with a module type by crevice depth interaction term to test depth effects within each module type. Planar recruit density was analyzed with a negative binomial GLM with log planar area as an offset.

## Results

### Effect of module design on coral recruitment

Across the 89 surveys at the four sites at 2 months, 6 months, and 1 year after spawning, recruit counts were highly variable (Fig. 3b). On average, 15.8 recruits were found per dome, with up to 381 recruits recorded on a Superdome at Hinalea Reef 2 months after spawning (Table S2). Recruitment patterns varied with site and shifted over time. At 2 months, counts were highest at Hinalea Reef and near zero at Lighthouse and Ulupaʻu. By 6 months, recruitment at Hinalea had declined while counts at Patch Reef 13 and Ulupaʻu increased. At 1 year, Patch Reef 13 recorded the highest counts overall, while Lighthouse and Ulupaʻu stabilized at around 2 recruits per structure.

After one year, dome type significantly affected both total recruit counts (Fig. 3c, NB GLMM, χ²(3) = 22.6, *p* < 0.001; Table S3) and surface-area-standardized recruit density (Fig. 3d, χ²(3) = 19.7, *p* < 0.001; Table S4). In absolute numbers, the Superdome attracted the most recruits and performed significantly better than all other designs (∼20× more than the Control). The Fractal Dome performed significantly better than the Control and Hexagon, but had fewer recruits than the Superdome (BH-adjusted pairwise contrasts; Table S3). After standardizing for surface area, only the Superdome had significantly more recruits per area unit than the Control and Hexagon (∼14× more), while the Fractal Dome was intermediate and no longer significantly different from any other design. Control and Hexagon Domes did not differ in either metric.

### Effect of reef environment on Superdome recruitment

Planar recruit density differed significantly among reef environments and the natural reef baseline (Fig. 3e, NB GLM, χ²(4) = 82.5, *p* < 0.001, Table S5). Domes at all sites recruited significantly more coral larvae than the natural reef. The highest density occurred at Patch Reef 13 (high cover, high flow; ∼32× reef baseline), followed by Hinalea Reef (medium cover, medium flow; ∼9×), Ulupaʻu (low cover, high flow; ∼4×), and Lighthouse Reef (medium cover, low flow; ∼3×).

### Recruitment performance of production-scale versus experimental Superdomes

The large concrete Superdome had the highest crevice occupancy rate (94%) and settlement probability differed significantly among module types (Fig. 4; binomial GLMM, χ²(2) = 9.49, *p* = 0.009; Table S6). Specifically, the large concrete Superdome had significantly higher settlement probabilities than the small concrete (50%; *p* = 0.028) but not small clay (75%; *p* = 0.162) Superdomes. Among occupied crevices, the large concrete Superdome showed lower recruit density per unit crevice area than both smaller module types (Gamma GLMM, χ²(2) = 18.1, *p* < 0.001; Table S6; BH-adjusted pairwise contrasts: vs. small concrete *p* = 0.040, vs. small clay *p* < 0.001). This difference was less pronounced when recruits were standardized to crevice perimeter length, where only the clay modules differed significantly from the large concrete Superdome (Gamma GLMM, χ²(2) = 13.1, *p* = 0.001), with no significant difference between the two concrete designs. Crevice depth had a significant effect on density standardized by crevice length (χ²(3) = 23.5, *p* < 0.001) and recess area (χ²(3) = 11.0, *p* = 0.012). For density standardized by crevice length, the effect was driven primarily by the small clay modules, whereas for density per recess area the only significant pairwise contrast was within the large concrete Superdome (depth 2 vs. 3; Table S7). In terms of planar recruit density, the large concrete Superdome performed comparably to the small concrete Superdome, but achieved approximately half the density of small clay (*p* < 0.001; NB GLM, χ²(2) = 24.4, *p* < 0.001; Table S8).

**Figure 4:**
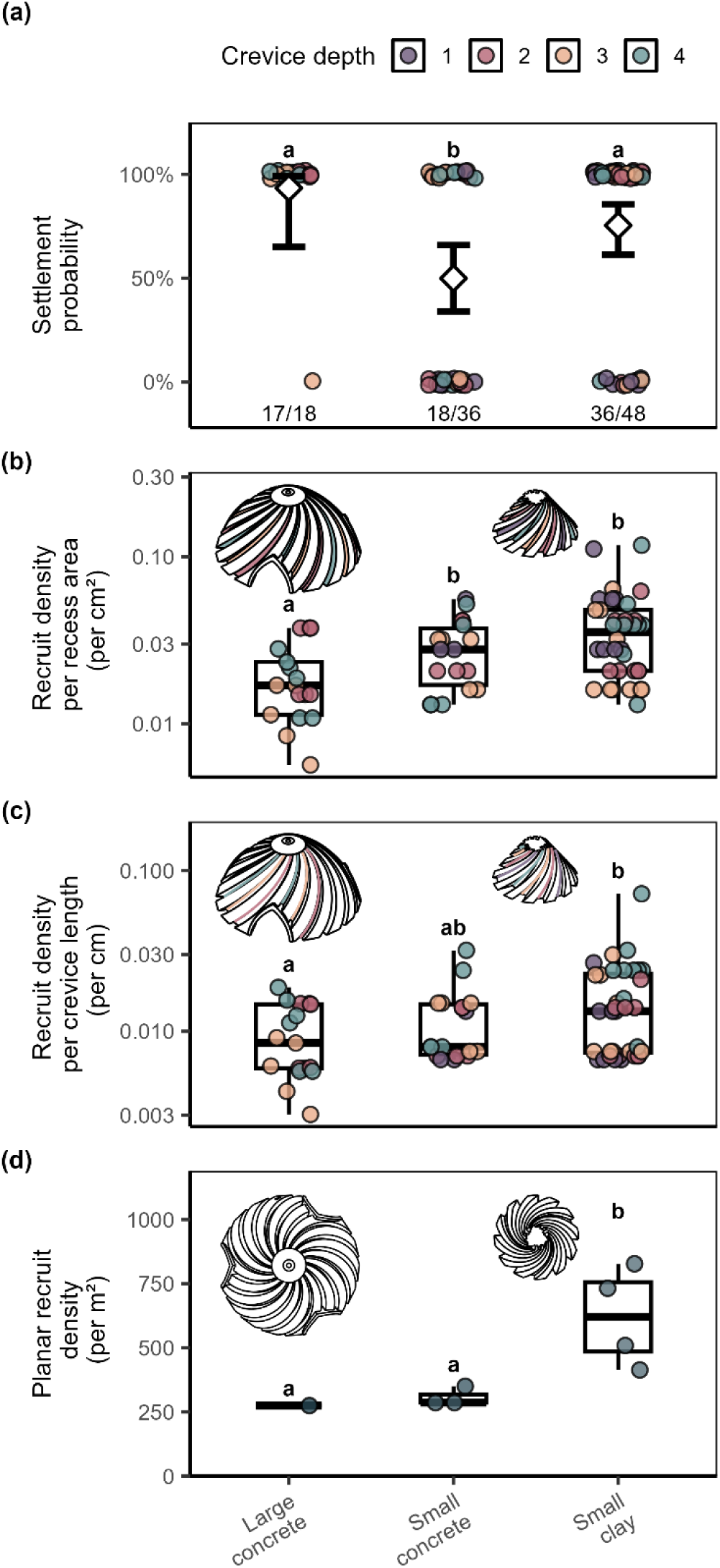
Coral recruit density on three Superdome module types (large concrete, small concrete, small clay) at Patch Reef 13. **(a)** Settlement probability (presence/absence) per module type; jittered points show raw binary observations, diamonds show estimated marginal means ± 95% CI from the binomial GLMM, and fractions below indicate observed n/N crevices occupied. **(b)** Recruit density per crevice recess area and **(c)** per crevice perimeter length, both among occupied crevices only. Individual points are colored by crevice depth category (1 = shallowest, 4 = deepest). **(d)** Planar recruit density per module planar surface area. Points represent individual modules. Letters denote significant differences from BH-adjusted pairwise emmeans contrasts of a hurdle GLMM with dome ID as a random effect (a–c) and a negative-binomial GLM (d).

### Large-scale module production

A total of 690 modules were produced between October 2024 and May 2026 (Fig. 5). No noticeable differences in surface quality or geometric fidelity were observed between modules produced at the precast facility and those produced during our initial study. However, some recurring manufacturing challenges were identified. (1) Thin concrete membranes frequently covered the openings of the Hexagon Dome holes and required manual removal. (2) The Hexagon Dome was also susceptible to cracking during transport, resulting in a structural failure rate of approximately 20% of the modules. (3) Fractal Domes were occasionally cast at a slight angle, causing the central attachment hole to exit off-center at the base, and some exhibited an uneven base due to concrete displacement during casting. (4) Exposed aggregate was occasionally visible on the top surface of all dome designs.

**Figure 5:**
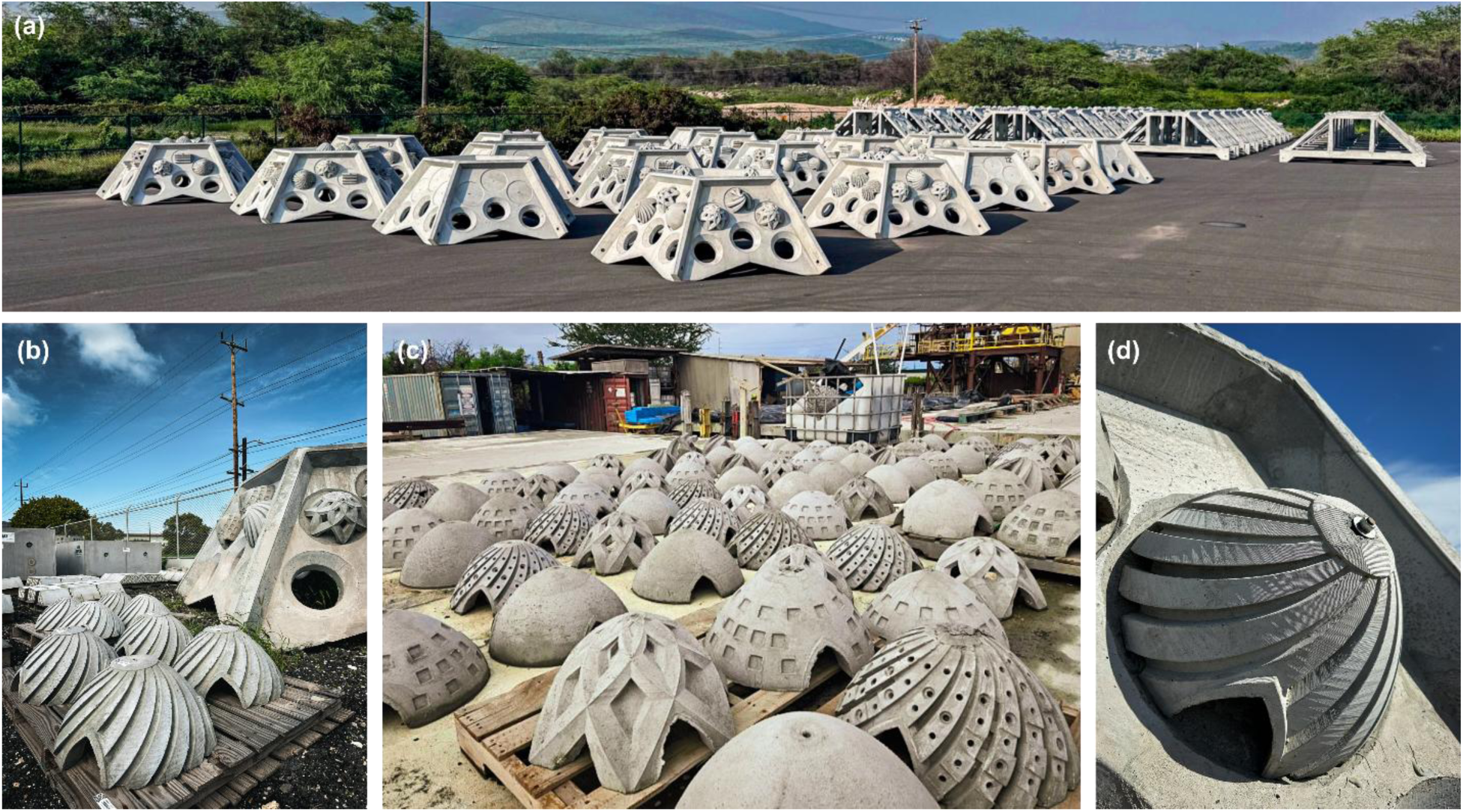
Large-scale production of the coral settlement modules for the R3D deployment. A total of 690 modules were produced from standardized molds at a precast concrete facility (Jensen Infrastructure, Kapolei, Oʻahu, Hawaiʻi) between October 2024 and May 2026. (a) Assembled hybrid reef structures (wave-breaker units) fitted with the settlement module designs, staged at the facility prior to deployment. (b) Cast concrete Superdomes with helix recesses on pallets, with an assembled reef structure in the background. (c) The range of cast module designs produced at volume, including Superdomes, Hexagon Domes, and Integration Domes (with insets) at the precast facility. (d) Detail of a cast Superdome showing the helix recesses and the central threaded-rod fitting used to anchor the module.

## Discussion

This study highlights the successful transfer of the helix-recess geometry from experimental clay modules to production-scale concrete structures, demonstrating consistent recruitment enhancement across four reef environments with contrasting coral cover and flow regimes. The module incorporating helix recesses outperformed the featureless control module approximately 20-fold, with the design hierarchy preserved regardless of local conditions. Production-scale concrete achieved settlement probability and planar recruit density comparable to small experimental concrete modules. A total of 690 modules were successfully produced for a large-scale deployment using standardized molds, demonstrating the feasibility of production at scale.

### Module design drives coral recruitment

Different module designs revealed varying success for coral recruitment. The Superdome attracted the most coral recruits, followed by the Fractal Dome, with the Hexagon and Control Domes performing equally poorly (Fig. 3). This pattern was most visible in absolute counts, although this trend also persisted after area standardization. This suggests that the advantage of the Fractal Dome is partly an area effect, attributable to its large surface area. The Superdome maintained its top ranking after standardization, indicating that its recess geometry provides per-area settlement benefits beyond surface area alone.

The Superdome’s consistent advantage is likely driven by flow dynamics and larval preference for cryptic, low-light microhabitats. Millimeter-scale topography within the recesses reduces near-surface velocities and extends larval contact time with the substrate (Levenstein, Gysbers, et al. 2022). Larvae preferentially settle in cryptic spaces at low light levels, balancing light availability with protection from predation and grazing (Doropoulos et al. 2016; Reichert et al. 2025). The design ranking was preserved across all four sites, extending the robustness of the helix recess effect to a wider environmental gradient.

### Superdome recruitment exceeds natural reef baseline across all environments

Production-scale Superdomes consistently outperformed adjacent natural reef substrate at all four sites. The fold excess ranged from ∼32× at Patch Reef 13 to ∼3× at Lighthouse Reef. These variations likely reflect the combined influence of local larval supply and flow regime. Coral cover is a direct proxy for adult spawner biomass, and reefs with greater adult abundance generally deliver higher larval concentrations to nearby substrates (Hughes et al. 2000). However, the rank order of recruitment across our sites cannot be explained by coral cover alone. Patch Reef 13 (high cover, high flow) produced the highest fold excess, consistent with abundant local larval supply combined with active flow delivery. The two medium-cover sites diverged sharply by flow regime: Hinalea Reef (medium flow) recruited at an intermediate level (∼9×), whereas Lighthouse Reef (low flow) showed the lowest fold excess (∼3×), suggesting that limited flow restricts larval transport from upstream sources. Ulupaʻu (low cover, high flow) recruited marginally more than Lighthouse (∼4× vs ∼3×) despite minimal local cover, indicating that flow-mediated import from adjacent reefs partly compensated for low local supply but chronic sedimentation at this site likely additionally suppresses settlement (Babcock & Davies 1991; Gilmour 1999). These findings are in accordance with other restoration studies. Site characteristics consistently drive more variation in the effectiveness of a restoration approach than device design (Ramsby et al. 2026; Waters et al. 2025), underscoring that pre-deployment site assessment is critical (Boström-Einarsson et al. 2020).

The fold-increase in recruit densities on structures compared to natural substrate observed here (3–32× planar density at 1 year) is in line with our previous studies on the experimental units. Coral settlement modules with helix recesses consistently outperformed adjacent natural reef substrate at 2 months, by ∼70-fold standardized to surface area (Reichert et al. 2025), and ∼300-fold standardized to planar area (Reichert et al. 2026). Given that these values differ in area standardization and census timepoint (2 months vs. 1 year), the lower increase observed here is likely attributed to the later census timepoint, at which post-settlement mortality might differently influence the fold-comparison. These results are comparable to other restoration approaches, targeting direct larval delivery, which report initial settlement enhancements of 40–305× above background (Gouezo et al. 2026; Doropoulos et al. 2025), depending on site conditions, post-settlement survival, and local larval supply (Edwards et al. 2015; Waters et al. 2025). However, the passive coral settlement approach, unlike active seeding approaches, intercepts naturally spawned larvae and can therefore sustain these enhancements across successive spawning events without additional effort (Reichert et al. unpublished data).

The lower end of this range warrants critical interpretation. Although the increases for the low-cover sites should be treated as conservative estimates since they were compared against the high-cover, high-flow Barrier Reef, a 3-fold enhancement over natural reef recruitment may be ecologically marginal for restoration purposes (Madin et al. 2025). At sites where the natural recruitment baseline is already low, low recruit densities are unlikely to shift population trajectories or accelerate meaningful recovery and passive enhancement cannot compensate for depleted larval pools. The effectiveness of passive recruitment structures depends on local larval supply, which can be influenced by factors such as large- and small-scale hydrodynamics, adult coral cover, reproductive output, and connectivity to adjacent source reefs. Deploying at sites without adequate larval supply is unlikely to yield ecologically meaningful recruitment gains, regardless of module design. Therefore, pre-deployment assessment of larval availability and habitat suitability should precede any large-scale deployment program. Small experimental coral settlement modules, which are easy to manufacture, deploy, retrieve, and relocate, are well-suited for this purpose (Reichert et al. 2025). Placing a subset of experimental units at candidate sites prior to full production commitment allows direct measurement of local recruitment potential under ambient conditions. Additional habitat suitability analyses incorporating substrate stability, sedimentation rates, water flow, and spatial connectivity with source reefs should accompany such assessments.

### Coral settlement modules maintain high recruitment success following production-scale translation

Despite the increase in module size, the production-scale Superdomes maintained high per-unit settlement performance. Planar recruit density on the large concrete Superdomes was similar to that of the small concrete modules (∼300 recruits m⁻²). Recruit density per crevice area was lower in the large modules than in both smaller types, reflecting the characteristics of recesses as settlement habitat for coral larvae. The fact that recruit density per crevice perimeter length was equivalent between large and small concrete modules indicates that the functional settlement habitat, the cryptic inner edge of the recess, was preserved at scale. Coral larvae preferentially settled along the cryptic inner edges of these recesses as observed in previous studies (Reichert et al. 2025). This indicates that edge length, not surface area, is the biologically relevant unit of available habitat. The recess dimensions of the production Superdomes may also benefit long-term post-settlement survival. Recruit density inside deeper recesses was higher, likely reflecting reduced competition and greater light availability within wider crevices as recruits grow (Reichert et al. 2026). Because coral recruits respond to absolute, millimeter-scale recess geometry rather than module size, the production Superdome retained the same absolute recess dimensions as the experimental modules, including the deeper recesses associated with the highest post-settlement survival, thereby preserving the proven microhabitat geometry at production scale. Together, these comparisons show that production-scale translation preserved the settlement performance of the experimental designs.

Material differences between concrete and clay were transient at short time scales but diverged over the long term. While our earlier study revealed a slight advantage of concrete over clay during the initial settlement phase (Reichert et al. 2026), the current study suggests a clay advantage after 1 year of deployment. The mechanism might be related to the higher surface pH of the concrete, which transiently delays marine colonization (Knoester et al. 2024), potentially causing a long-term colonization disadvantage for the concrete domes. The lower porosity of the concrete may further reduce long-term biofilm accumulation compared to fired clay, contributing to the observed divergence. However, because we did not measure surface pH or biofilm development, these proposed mechanisms remain hypotheses requiring further testing. Emerging concrete formulations designed for marine deployment offer routes to address these limitations (Dennis et al. 2018; Yus et al. 2024; Ruszczyk et al. 2026).

## Limitations

Several limitations of this study should be acknowledged. Recruits were monitored to 1 year post-spawning, at which point they remain small juveniles. Multi-year monitoring will better assess long-term demographic impacts. Recruit species identity was not determined, since the snorkel-based census did not allow for discrimination among taxa, leaving the relative contributions of species unresolved. Additionally, because the census relied on fluorescence under blue light, non-fluorescing recruits may have gone undetected, meaning reported recruitment rates might be underestimated (Baird et al. 2006). Also, the replication was limited, especially at two sites (n = 1 Superdome at Patch Reef 13; n = 1 Fractal at Lighthouse Reef), and estimates from these sites carry greater uncertainty. Further, testing this approach on reefs beyond Oʻahu, with different coral communities, hydrodynamics, and sedimentation regimes, is necessary to establish its broader applicability.

Among established larval restoration approaches, the passive coral settlement module is unique in that it requires no rearing infrastructure, no larval handling, and no fragment outplanting. Active seeding approaches achieve higher initial settlement densities and allow targeted delivery of genetically diverse stocks, but require substantial investment in larval culture, transport, and deployment logistics. We conclude that passive structures are therefore a scalable complement to active restoration measures that warrant further investigation on larger scales.

### Feasibility and recommendations for mass production of coral settlement modules

Large-scale concrete casting of geometrically precise settlement modules proved logistically feasible. A total of 690 modules were produced for the R3D wave breaker deployment (DARPA 2024), without loss of helix recess fidelity across units. Casting in standardized molds allowed rapid production at a fraction of the time and cost of equivalent-volume 3D-printed ceramic modules. Production success varied across designs, with recurring challenges specific to the Hexagon and Fractal Dome designs. Approximately 20% of the Hexagon modules failed due to cracking at the top during demolding or curing. Fractal Domes presented geometric alignment challenges due to the lack of the inner cave, which increased production time per unit relative to the Superdome and Control. These challenges are addressable through mold redesign and do not represent fundamental constraints on scaling. The Superdome, combining the highest recruitment performance with the most straightforward casting process, is the most production-ready design for immediate upscaling.

The Hexagon design had the fewest coral recruits, performed equivalently to the featureless control, and is not recommended as a primary structure when the goal is to increase coral recruitment. However, three-dimensional rugosity provides value beyond coral recruitment. In Kāneʻohe Bay specifically, this module design was used as a fish habitat module and attracted larval fish in the scope of an acoustic enrichment study (Boulais et al. 2023, 2026). Structural complexity is a key driver of fish community structure more broadly. Across the Main Hawaiian Islands, reef rugosity was among the strongest predictors of diverse fish assemblages (Asbury et al. 2026), and artificial reefs most reliably benefit fish communities as nursery habitat (Higgins et al. 2022). Including a high-rugosity design in a mixed array therefore contributes structural complexity that benefits fish and invertebrate assemblages at reef scale.

### Integration with emerging restoration technologies

The passive coral settlement module design is compatible with a range of emerging restoration technologies, several of which are being tested in dedicated integration modules within the large-scale deployment. An integration of plugholes allows for the attachment of coral fragments directly onto the structure, enabling outplanting of fragments of opportunity or thermally tolerant genotypes alongside passive larval recruitment (Hoʻopai-Sylva et al. 2026). Settlement-inducing surface treatments such as the nanoparticle gel releasing crustose coralline algae-derived chemical cues SNAP-X, can be applied to indents on recess surfaces and ridges to enhance larval capture (Kundu et al. 2025). Similarly, millimeter-scale bioinspired microtextures can be attached to or cast into the module surfaces to create hydrodynamic and low-light refugia, additionally increasing larval settlement (Levy et al. in review). Emerging technologies, like Underwater Zooplankton Enhancement Light Array (UZELA) devices, can be housed within the dome cavity to attract zooplankton at night, increasing heterotrophic prey availability for recruits in shaded recess microhabitats (Grottoli et al. 2025; Dixon et al. 2025, Jorissen et al. in review). Acoustic enrichment speakers within the cavity broadcast healthy reef soundscapes to attract settlement-stage fish larvae, with playback shown to increase larval fish presence (Boulais et al. 2023, 2026; Thode et al. in review). These integration modules demonstrate that passive settlement structures can serve as a platform for combining multiple restoration approaches within a single deployed unit. Beyond technological integration, the modular design itself supports broad geographic accessibility. Silicone molds can be shipped to remote locations and modules cast using locally available concrete, removing dependence on specialized materials or supply chains. This makes the approach well-suited to cost-efficient, community-led restoration initiatives that can be implemented across diverse reef regions worldwide.

Furthermore, the modules themselves offer a range of application potential that extends beyond conventional stand-alone reef restoration. The most direct application is the placement of arrays on degraded reef substrate to passively enhance natural recruitment. A second avenue is the integration into new or existing coastal infrastructure (DARPA 2024). A third application is their use as substrate consolidation mattresses on rubble-dominated reef zones. Interlocking or tethered module configurations could stabilize loose substrate while simultaneously providing enhanced settlement habitat, addressing two barriers to reef recovery at once.

We recommend deploying mixed-design arrays. The helix-recess module should be the primary structure for targeting coral recruitment, given its consistent performance and straightforward casting process. A multi-level complexity design like the Fractal Dome is recommended as a secondary component to broaden taxonomic diversity of recruits. Designs like the Hexagon Dome contribute mainly to structural complexity for fish and invertebrate assemblages and serve best as a supplementary element for overall reef biodiversity. Deployments should prioritize sites with adequate larval supply and consolidated substrate, validated through pre-deployment ecological assessment. Future work should test the approach across additional reef systems and coral communities to confirm generality. Given the effectiveness of the helix recess design and its demonstrated scalability for large-scale production, passive seeding approaches based on coral settlement modules have considerable potential to enhance the logistical and economic feasibility of reef restoration at ecologically meaningful scales.

## Acknowledgements

We thank members of the Marine Conservation Innovation Group at the Hawaiʻi Institute of Marine Biology, who supported the production and fieldwork of this study. This work was funded by the Defense Advanced Research Projects Agency (Contract No. HR001122C0134), the National Science Foundation (1948946), and the HIMB Director’s Innovation Fund.

## Author contributions

Conceptualization: JR, RA, BJ, JL, AT, DW, JM

Data Curation: JR

Formal Analysis: JR

Funding Acquisition: RA, BJ, AT, DW, JM

Investigation: JR, MA, GKC, JE, HJ, JL, ADN, JM

Methodology: JR, LR, MR, JM

Project Administration: JR, BJ, JL, JM

Resources: RA, BJ, JL, AT, DW, JM

Visualization: JR

Writing – Original Draft Preparation: JR

Writing – Review & Editing: All authors

## Data Availability Statement

The data and R code supporting this study are available at https://github.com/JessiReichert/Coral_settlement_modules upon acceptance.

## Use of Generative AI

Claude (Anthropic PBC, San Francisco, CA, USA) was used to assist with manuscript editing, including reorganizing text, reducing word count, and checking consistency across sections, and to assist with R code development for statistical analyses. All scientific content, data interpretation, and conclusions are solely the authors’ own. The authors take full responsibility for the accuracy and integrity of the work.

## References

Asbury M, Schiettekatte NMD, Kindinger TL, Richardson L, Madin JS (2026) Benthic habitat structure explains broad-scale patterns in reef fish communities. Ecological Applications 36(3):e70238.

Babcock R, Davies P (1991) Effects of sedimentation on settlement of *Acropora millepora*. Coral Reefs 9:205–208.

Baird AH, Salih A, Trevor-Jones A (2006) Fluorescence census techniques for the early detection of coral recruits. Pages 73–76. Coral Reefs. Vol. 25

Bates D, Mächler M, Bolker BM, Walker SC, Machler M, Bolker BM, Walker SC (2015) Fitting Linear Mixed-Effects Models Using lme4. Journal of Statistical Software 67:1–48.

Boström-Einarsson L, Babcock RC, Bayraktarov E, Ceccarelli D, Cook N, Ferse SCA, Hancock B, Harrison P, Hein M, Shaver E, Smith A, Suggett D, Stewart-Sinclair PJ, Vardi T, McLeod IM (2020) Coral restoration – A systematic review of current methods, successes, failures and future directions. PLoS ONE 15.

Boulais O, Schar D, Levy J, Kim K, Levy N, Reichert J, Schiettekatte N, Wangpraseurt D, Madin J, Thode AM (2023) Acoustic enrichment trials using autonomous cameras on a Hawaiian coral reef. The Journal of the Acoustical Society of America 154:A22–A22.

Boulais O, Thode A, Pickering C, Levy N, Schar D, Reichert J, Maldonado T, Madin JS, Levy J, the R3D Consortium (2026) Autonomous cameras reveal larval reef fish responses to acoustic enrichment and lunar phase. Scientific Reports (accepted)

Brooks ME, Kristensen K, Van Benthem KJ, Magnusson A, Berg CW, Nielsen A, Skaug HJ, Mächler M, Bolker BM (2017) glmmTMB Balances Speed and Flexibility Among Packages for Zero-inflated Generalized Linear Mixed Modeling.

Carpenter KE, Abrar M, Aeby G, Aronson RB, Banks S, Bruckner A, Chiriboga A, Cortes J, Delbeek JC, DeVantier L, Edgar GJ, Edwards AJ, Fenner D, Guzman HM, Hoeksema BW, Hodgson G, Johan O, Licuanan WY, Livingstone SR, Lovell ER, Moore JA, Obura DO, Ochavillo D, Polidoro BA, Precht WF, Quibilan MC, Reboton C, Richards ZT, Rogers AD, Sanciangco J, Sheppard A, Sheppard C, Smith J, Stuart S, Turak E, Veron JEN, Wallace C, Weil E, Wood E (2008) One-Third of Reef-Building Corals Face Elevated Extinction Risk from Climate Change and Local Impacts. Science 321:560–563.

Ceccarelli DM, McLeod IM, Bostrom-Einarsson L, Bryan SE, Chartrand KM, Emslie MJ, Gibbs MT, Rivero MG, Hein MY, Heyward A, Kenyon TM, Lewis BM, Mattocks N, Newlands M, Schlappy ML, Suggett DJ, Bay LK (2020) Substrate stabilisation and small structures in coral restoration: State of knowledge, and considerations for management and implementation. PLoS ONE 15.

Chamberland VF, Petersen D, Guest JR, Petersen U, Brittsan M, Vermeij MJA (2017) New Seeding Approach Reduces Costs and Time to Outplant Sexually Propagated Corals for Reef Restoration. Scientific Reports 7.

Cruz DWD, Harrison PL (2017) Enhanced larval supply and recruitment can replenish reef corals on degraded reefs. Scientific Reports 7.

DARPA (2024) Reefense program. Defense Advanced Research Projects Agency. https://www.darpa.mil/research/programs/reefense (accessed 5 June 2026)

Dennis HD, Evans AJ, Banner AJ, Moore PJ (2018) Reefcrete: Reducing the environmental footprint of concretes for eco-engineering marine structures. Ecological Engineering 120:668–678.

Dixon SL, Jorissen H, Hulver AM, Welter J, Toonen RJ, the R3D Consortium, Madin JS, Grottoli AG (2025) Technology Solutions for Overcoming the Coral Recruitment Bottleneck. Environmental Science and Technology 59:16368–16378.

Doropoulos C, Roff G, Bozec YM, Zupan M, Werminghausen J, Mumby PJ (2016) Characterizing the ecological trade-offs throughout the early ontogeny of coral recruitment. Ecological Monographs 86:20–44.

Doropoulos C, Roff G, Carlin G, Gouezo M, dela Cruz D, Chai A, Hardiman L, Hasson L, Thomson DP, Harrison PL (2025) Larval seedboxes: A modular and effective tool for scaling coral reef restoration. Ecological Applications 35.

Edmunds PJ (2023) Coral recruitment: patterns and processes determining the dynamics of coral populations. Biological Reviews 98:1862–1886.

Edwards AJ, Guest JR, Heyward AJ, Villanueva RD, Baria MV, Bollozos ISF, Golbuu Y (2015) Direct seeding of mass-cultured coral larvae is not an effective option for reef rehabilitation. Marine Ecology Progress Series 525:105–116.

Gilmour J (1999) Experimental investigation into the effects of suspended sediment on fertilisation, larval survival and settlement in a scleractinian coral. Marine Biology 135:451–462.

Gouezo M, Harrison PL, Roff G, Chai A, Thomson DP, Guglielmo M, Hardiman L, Forbes A, Gardner B, Doropoulos C (2026) Coral larval enhancement with and without nets yields similar recruitment during slack current releases. Restoration Ecology 34.

Grottoli AG, Dixon SL, Hulver AM, Bardin CE, Lewis CJ, Suchocki CR, Toonen RJ (2025) Underwater Zooplankton Enhancement Light Array (UZELA): A technology solution to enhance zooplankton abundance and coral feeding in bleached and non-bleached corals. Limnology and Oceanography: Methods 23:201–211.

Hall VR, Hughes TP (1996) Reproductive Strategies of Modular Organisms: Comparative Studies of Reef-Building Corals. Ecology 77:950–963.

Harrison PL, dela Cruz DW, Cameron KA, Cabaitan PC (2021) Increased Coral Larval Supply Enhances Recruitment for Coral and Fish Habitat Restoration. Frontiers in Marine Science 8.

Harrison PL, Wallace CC (1990) Reproduction, dispersal and recruitment of scleractinian corals. Ecosystems of the world 25:133–207.

Hata T, Madin JS, Cumbo VR, Denny M, Figueiredo J, Harii S, Thomas CJ, Baird AH (2017) Coral larvae are poor swimmers and require fine-scale reef structure to settle. Scientific Reports 7.

Higgins E, Metaxas A, Scheibling RE (2022) A systematic review of artificial reefs as platforms for coral reef research and conservation. Chapman, M (Gee) G (ed.) PLOS ONE 17:e0261964.

Hoʻopai-Sylva H, Caruso C, Miller S, Hancock JR, Parry M, Hughes K, Drury C (2026) Proactive Coral Reef Restoration Using Thermally Tolerant Corals in Hawaiʻi. Conservation Letters 19.

Hughes TP, Baird AH, Dinsdale EA, Moltschaniwskyj NA, Pratchett MS, Tanner JE, Willis BL (2000) Supply-side ecology works both ways: the link between benthic adults, fecundity, and larval recruits. Ecology 81(8):2241–2249.

Hughes TP, Barnes ML, Bellwood DR, Cinner JE, Cumming GS, Jackson JBC, Kleypas J, Van De Leemput IA, Lough JM, Morrison TH, Palumbi SR, Van Nes EH, Scheffer M (2017) Coral reefs in the Anthropocene. Nature 546:82–90.

IPCC (2023) Climate Change 2023: Synthesis Report. Contribution of Working Groups I, II and III to the Sixth Assessment Report of the Intergovernmental Panel on Climate Change [Core Writing Team, H. Lee and J. Romero (eds.)]. IPCC, Geneva, Switzerland.

Jorissen H, Galand PE, Bonnard I, Meiling S, Raviglione D, Meistertzheim AL, Hédouin L, Banaigs B, Payri CE, Nugues MM (2021) Coral larval settlement preferences linked to crustose coralline algae with distinct chemical and microbial signatures. Scientific Reports 11.

Jorissen H, Reichert J, Dixon SL, Madin EM, Madin CA, Asbury M, Nims AD, Rottmueller ME, Rova L, Grottoli AG, the R3D Consortium, Madin JS (in review) Field testing the influence of settlement recesses and nighttime feeding lights on coral recruitment.

Knoester EG, Vos A, Saru C, Murk AJ, Osinga R (2024) Concrete evidence: outplanted corals for reef restoration do not need extended curing of ordinary Portland cement. Royal Society Open Science 11.

Kundu S, Potenti S, Quinlan ZA, Willard H, Chen J, Noritake T, Levy N, Karimi Z, Jorissen H, Hancock JR, Drury C, Kelly LW, De Cola L, Chen S, Wangpraseurt D (2025) Biomimetic chemical microhabitats enhance coral settlement. Trends in Biotechnology 43:2232–2250.

Lenth R V., Piaskowski J, Banfai B, Bolker B, Buerkner P, Giné-Vázquez I, Hervé M, Jung M, Love J, Miguez F, Riebl H, Singmann H (2025) emmeans: Estimated Marginal Means, aka Least-Squares Means. R package.

Leung SK, Kenyon TM, Raymundo LJ, Fox HE, Cook N, Cook K, Edwards AJ, Fisher EE, Brival AJT, Nicholson FE, Philippo RWL, Taylor ACF, Bryan SE, Lewis BM, Abdul Adzis KA Bin, Edmondson JP, Griffin SP, Li X, Liu X, Oakley HA, Razak TB, Samudra SH, Welly M, Mumby PJ (2025) A decision support tool for rubble stabilization on coral reefs. Journal of Environmental Management 396.

Levenstein MA, Gysbers DJ, Marhaver KL, Kattom S, Tichy L, Quinlan Z, Tholen HM, Kelly LW, Vermeij MJA, Wagoner Johnson AJ, Juarez G (2022) Millimeter-scale topography facilitates coral larval settlement in wave-driven oscillatory flow. PLoS ONE 17.

Levenstein MA, Marhaver KL, Quinlan ZA, Tholen HM, Tichy L, Yus J, Lightcap I, Wegley Kelly L, Juarez G, Vermeij MJA, Wagoner Johnson AJ (2022) Composite Substrates Reveal Inorganic Material Cues for Coral Larval Settlement. ACS Sustainable Chemistry and Engineering 10:3960–3971.

Levy N, Berman O, Yuval M, Loya Y, Treibitz T, Tarazi E, Levy O (2022) Emerging 3D technologies for future reformation of coral reefs: Enhancing biodiversity using biomimetic structures based on designs by nature. Science of The Total Environment 830:154749.

Levy N, Pimenta P, Inman B, Alvaro A, Smith Z, Kundu S, Karimi Z, Teng Chua S, Dhar N, Galindo-Martínez CT, Reichert J, Madin J, Drury C, Hancock J, Chen S, Jaffe J, Kurtis Cole M, Levy J, Verma S, the R3D Consortium, Wangpraseurt D (In review) Engineering Microhabitats to Enhance Coral Larval Settlement.

Lowe RJ, Falter JL, Monismith SG, Atkinson MJ (2009) A numerical study of circulation in a coastal reef-lagoon system. Journal of Geophysical Research: Oceans 114.

Madin JS, Oliver T, McWilliam M, Asbury M, Baird AH, Chen GK, Connolly SR, Couch C, Drury C, Ehrenberg J, Jorissen H, Nims AD, Reichert J, Rova L, Schiettekatte NMD, Toonen RJ, Wulstein DM (2025) Demographic insights for coral restoration. bioRxiv. 10.1101/2025.10.21.683544

Nozawa Y (2008) Micro-crevice structure enhances coral spat survivorship. Journal of Experimental Marine Biology and Ecology 367:127–130.

Pizarro O, Friedman A, Bryson M, Williams SB, Madin J (2017) A simple, fast, and repeatable survey method for underwater visual 3D benthic mapping and monitoring. Ecology and Evolution 7:1770–1782.

R Core Team (2025) R: A language and environment for statistical computing. http://www.r-project.org/

Ramsby BD, Forster R, Ferguson SN, Haikola P, Randall CJ, Abdul Wahab MA, Mead DJ, Severati A (2026) Developing coral seeding devices and rapid deployment methods to scale up reef restoration. Restoration Ecology 34.

Randall C, Negri A, Quigley K, Foster T, Ricardo G, Webster N, Bay L, Harrison P, Babcock R, Heyward A (2020) Sexual production of corals for reef restoration in the Anthropocene. Marine Ecology Progress Series 635:203–232.

Reichert J, Jorissen H, Drury C, Hancock JR, Haynes C, Nims AD, Rova LH, Schiettekatte NMD, Madin JS (2025) Helix recesses boost coral larvae settlement and survival. Biological Conservation 311.

Reichert J, Jorissen H, Rottmueller ME, Nims AD, Rova LH, Drury C, Madin JS (2026) Cylindrical, pylon-like structures with helix recesses enhance coral larval recruitment. Ecological Engineering 226:107912.

Roach TNF, Yadav S, Caruso C, Dilworth J, Foley CM, Hancock JR, Huckeba J, Huffmyer AS, Hughes K, Kahkejian VA, Madin EMP, Matsuda SB, McWilliam M, Miller S, Santoro EP, Rocha de Souza M, Torres-Pullizaa D, Drury C, Madin JS (2021) A Field Primer for Monitoring Benthic Ecosystems using Structure-from-Motion Photogrammetry. Journal of Visualized Experiments 1–14.

Ruszczyk M, Rodriguez S, Tuen M, Rux K, Paul V, Chandragiri S, Stickley M, Swart PK, Haus BK, Baker AC, Miller MW, Suraneni P, Langdon C, Prakash VN (2026) Alkalinity-enhanced artificial substrates modulate local pH and increase survivorship of early-stage coral recruits. Communications Earth and Environment 7.

Schiettekatte NMD, Crichton R, Rova LH, Nims AD, Haynes C, Nishida T, Chapeau M, Chen GK, Cant J, Caruso C, Drury C, Hancock JR, Jorissen H, Madin EMP, Reichert J, Rutkowski E, Suzuki H, Suchocki CR, Toonen RJ, Vicente J, Madin JS (in review) Coral recruitment and biodiversity on the upward spiral.

Suggett DJ, van Oppen MJH (2022) Horizon scan of rapidly advancing coral restoration approaches for 21st century reef management. Emerging Topics in Life Sciences 6:125–136.

Thode A, Levy N, Boulais O, Kundu S, Freckelton M, Schar D, Hadfield M, Reichert J, Jorissen H, Kramer N, Levy J, Hughes K, the R3D Consortium, Madin J, Wangpraseurt D (in review) In situ enhancement of coral larval settlement using acoustic enrichment, engineered habitats and living materials.

Waters C, Harrison PL, Gouezo M, Severati A, Doropoulos C (2025) Early-stage coral survivorship using wild larval assemblages on coral seeding devices for reef restoration. Restoration Ecology 33.

Whalan S, Abdul Wahab MA, Sprungala S, Poole AJ, De Nys R (2015) Larval settlement: The role of surface topography for sessile coral reef invertebrates. PLoS ONE 10.

Yus J, Nixon EN, Li J, Noriega Gimenez J, Bennett MJ, Flores D, Marhaver KL, Wegley Kelly L, Espinosa-Marzal RM, Wagoner Johnson AJ (2024) Composite substrates for coral larval settlement and reef restoration based on natural hydraulic lime and inorganic strontium and magnesium compounds. Ecological Engineering 202.

