## Supplementary material for "Coral settlement module designs for scalable reef restoration": Figure S1

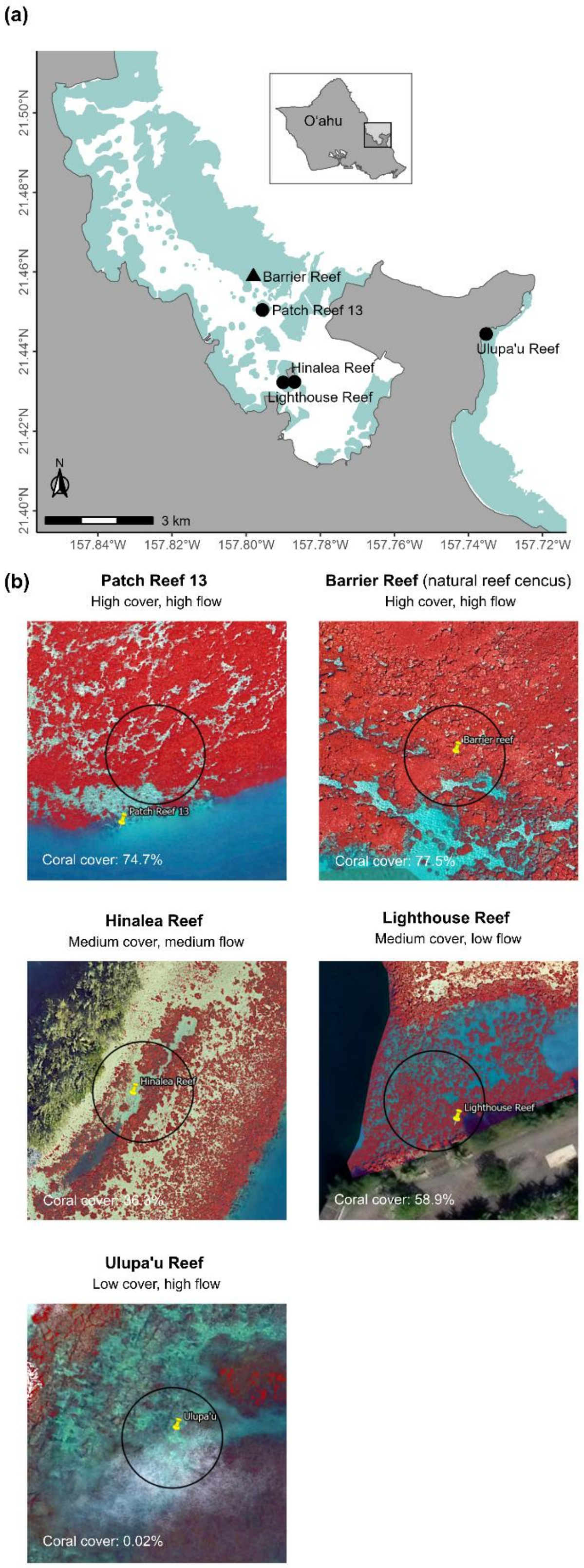


Figure S1: Coral cover at the study locations. (a) Map of the study locations in Kāneʻohe Bay on the windward side of Oʻahu, Hawaiʻi. Coral settlement modules were deployed at four reef sites (circles: Patch Reef 13, Hinalea Reef, Lighthouse Reef, and Ulupaʻu Reef), and natural reef recruitment was assessed at the Barrier Reef (triangle). Light green areas indicate the presence of reef. The inset shows the position of the study area on Oʻahu. (b) Google Earth satellite and aerial imagery of each location used to estimate coral cover. Coral cover was estimated within a 30m diameter area around each site (black circle) using the Image Classification tools in ArcGIS Pro 3.6.1, where an ISO Cluster unsupervised classifier was first used to identify 15 classes, followed by manual visual screening to select the classes corresponding to coral reef cover, with classified coral shown in red and the yellow marker indicating each site. Labels give the assigned cover and flow regime category and the estimated coral cover for each location (Patch Reef 13: 74.7%; Barrier Reef: 77.5%; Hinalea Reef: 36.3%; Lighthouse Reef: 58.9%; Ulupaʻu Reef: 0.02%). Sites were categorized as high (≥60%), medium (20–60%), or low (<20%) cover. These satellite-derived values are coarse, descriptive estimates used only to assign sites to cover categories and were not included in any statistical analysis.

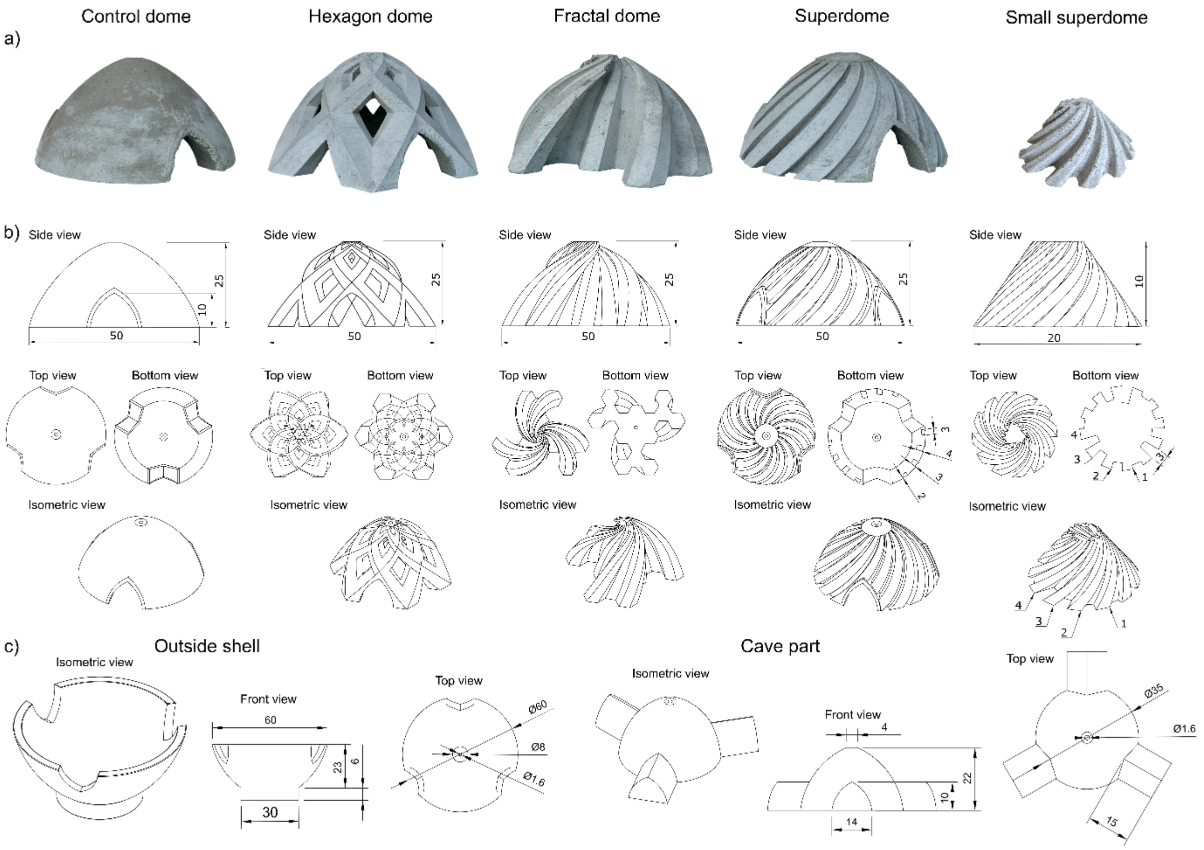


Figure S2: Coral settlement module designs and casting mold components. (a) Photographs of the five module designs: four large concrete modules (Control Dome, Hexagon Dome, Fractal Dome, Superdome; 50 cm diameter, 25 cm height) and the Small Superdome (20 cm diameter, 10 cm height), produced in clay and concrete as the size- and material-matched experimental comparison. (b) Technical drawings of each module (side, top, bottom, and isometric views; dimensions in cm). Bottom and isometric views of the Superdome and Small Superdome indicate the helix-recess depth categories (Superdome: three categories, numbered 2–4; Small Superdome: four categories, numbered 1 = shallowest to 4 = deepest), the shallowest category (1) having been omitted from the production Superdome. (c) Technical drawings of the casting mold components for the large modules, showing the outside shell and the internal cave part (isometric, front, and top views; dimensions in cm), which were mounted together using a threaded rod through the central Ø1.6 cm channel to align the mold and form.

Table S1: Properties of coral settlement modules. The shallowest crevice category previously tested in the Small Superdome (10 mm), was excluded during upscaling because it consistently yielded the lowest recruit density in previous work. All modules except the Fractal Dome incorporated an internal cavity to reduce weight and provide shelter for fish. A central attachment hole in the top of each module allowed anchoring to structures or the seafloor using a threaded rod. The faceted design of the Fractal Dome precluded the inclusion of an internal cavity.

| Module design | Key structural features | Recess dimensions | Surface area ($cm^2$) | Weight (kg) | Internal cave |
|---|---|---|---|---|---|
| Control Dome | Smooth dome | None | 3,071 | 27.1 | Yes |
| Hexagon Dome | 15 openings of 3 sizes | Small: 19 × 36 mm; Medium: 80 × 84 mm; Large: 110 × 75 mm | 4,125 | 14 | Yes |
| Fractal Dome | Three shelter levels (exposed, sheltered, cryptic) | Recesses: 47 mm deep × 88 mm wide at base, tapering to 5 mm deep × 14 mm wide at top | 4,579 | 22.2 | No |
| Superdome | 18 helix recesses with 3 depth categories | Depths: 20, 30, and 40 mm at base tapering to 2.5, 4, and 6 mm at top<br>Widths: 30 mm at base tapering to 7 mm at top | 4,458 | 18.1 | Yes |
| Small Superdome | 12 helix recesses with 4 depth categories | Depths: 10, 20, 30, and 40 mm at base tapering to 2, 5, 8, and 11 mm at top<br>Widths: 30 mm at base tapering to 6 mm at top | 306 | 1 | Yes |

Table S2: Mean total coral recruits per structure by location and timepoint.

| Location | Timepoint | n | Mean | SD | Median | Max |
|---|---|---|---|---|---|---|
| Hinalea Reef | 2 months | 3 | 340.7 | 36.8 | 332 | 381 |
| | 6 months | 3 | 36 | 21.6 | 30 | 60 |
| | 1 year | 3 | 15.3 | 4.2 | 14 | 20 |
| Lighthouse | 2 months | 10 | 0 | 0 | 0 | 0 |
| | 6 months | 10 | 0 | 0 | 0 | 0 |
| | 1 year | 10 | 1.9 | 2.6 | 0 | 7 |
| Patch Reef 13 | 2 months | NA | NA | NA | NA | NA |
| | 6 months | 1 | 62 | NA | 62 | 62 |
| | 1 year | 1 | 54 | NA | 54 | 54 |
| Ulupa‘u | 2 months | 16 | 0.6 | 1.1 | 0 | 4 |
| | 6 months | 16 | 3.1 | 4.7 | 2 | 17 |
| | 1 year | 16 | 2.1 | 4.3 | 0 | 17 |

Table S3: Pairwise contrasts from the negative binomial generalized linear mixed model for total recruit counts at 1 year (Lighthouse and Ulupaʻu reefs), with dome identity nested within reef location as random intercepts. Ratio = count ratio (first / second group). p-values adjusted across all dome-type contrasts using the Benjamini-Hochberg procedure. Bold p-values indicate statistical significance.

| Contrast | | | Ratio | SE | z | p (BH) |
|---|---|---|---|---|---|---|
| Control Dome | — | Hexagon Dome | 1.94 | 2.517 | 0.51 | 0.6070 |
| Control Dome | — | Fractal Dome | 0.16 | 0.143 | -2.04 | **0.0499** |
| Control Dome | — | Superdome | 0.05 | 0.04 | -3.62 | **0.0018** |
| Hexagon Dome | — | Fractal Dome | 0.08 | 0.093 | -2.19 | **0.0499** |
| Hexagon Dome | — | Superdome | 0.02 | 0.027 | -3.40 | **0.0021** |
| Fractal Dome | — | Superdome | 0.3 | 0.176 | -2.05 | **0.0499** |

Table S4: Pairwise contrasts from the negative binomial generalized linear mixed model for recruit densities at 1 year (Lighthouse and Ulupaʻu reefs), with dome identity nested within reef location as random intercepts. Ratio = count ratio (first / second group). p-values adjusted across all dome-type contrasts using the Benjamini-Hochberg procedure. Bold p-values indicate statistical significance.

| Contrast | | | Ratio | SE | z | p (BH) |
|---|---|---|---|---|---|---|
| Control Dome | — | Hexagon Dome | 2.61 | 3.3803 | 0.74 | 0.4580 |
| Control Dome | — | Fractal Dome | 0.23 | 0.2131 | -1.60 | 0.1320 |
| Control Dome | — | Superdome | 0.07 | 0.0581 | -3.18 | **0.0045** |
| Hexagon Dome | — | Fractal Dome | 0.09 | 0.1031 | -2.10 | 0.0537 |
| Hexagon Dome | — | Superdome | 0.03 | 0.0289 | -3.32 | **0.0045** |
| Fractal Dome | — | Superdome | 0.29 | 0.1714 | -2.10 | 0.0537 |

Table S5: Pairwise contrasts from the negative binomial generalized linear model comparing planar recruit density (per m²) among large concrete Superdome habitats and natural reef substrate at 1 year. Ratio = density ratio (first / second group). p-values adjusted using the Benjamini-Hochberg procedure. Bold p-values indicate statistical significance.

| Contrast | | | Ratio | SE | p (BH) |
|---|---|---|---|---|---|
| Patch Reef 13 | — | Hinalea Reef | 3.52 | 1.78 | **0.0159** |
| Patch Reef 13 | — | Lighthouse Reef | 10.8 | 5.92 | **<0.0001** |
| Patch Reef 13 | — | Ulupaʻu Reef | 8.31 | 4.23 | **<0.0001** |
| Patch Reef 13 | — | Natural reef | 31.73 | 14.7 | **<0.0001** |
| Hinalea Reef | — | Lighthouse Reef | 3.07 | 1.36 | **0.0159** |
| Hinalea Reef | — | Ulupaʻu Reef | 2.36 | 0.93 | **0.0322** |
| Hinalea Reef | — | Natural reef | 9.01 | 2.99 | **<0.0001** |
| Lighthouse Reef | — | Ulupaʻu Reef | 0.77 | 0.34 | 0.5570 |
| Lighthouse Reef | — | Natural reef | 2.94 | 1.16 | **0.0103** |
| Ulupaʻu Reef | — | Natural reef | 3.82 | 1.29 | **0.0001** |

Table S6: Pairwise contrasts from the hurdle generalized linear mixed model for crevice-level recruitment at Patch Reef 13. Binomial component (settlement probability) contrasts are on the odds-ratio scale. Gamma component (occupied crevices only) contrasts are on the response ratio scale. p-values adjusted across all dome-type contrasts using the Benjamini-Hochberg procedure. Bold p-values indicate statistical significance.

| Contrast | | | Ratio | SE | z | p (BH) |
|---|---|---|---|---|---|---|
| Binomial: settlement probability (odds ratios) | | | | | | |
| Large concrete | — | Small concrete | 14.24 | 15.52 | 2.44 | **0.0281** |
| Large concrete | — | Small clay | 4.62 | 5.06 | 1.40 | 0.1620 |
| Small concrete | — | Small clay | 0.33 | 0.16 | -2.35 | **0.0281** |
| Gamma: density per recess area (per $cm^2$, occupied crevices) | | | | | | |
| Large concrete | — | Small concrete | 0.69 | 0.12 | -2.22 | **0.0401** |
| Large concrete | — | Small clay | 0.53 | 0.08 | -4.23 | **0.0001** |
| Small concrete | — | Small clay | 0.77 | 0.11 | -1.85 | 0.0639 |
| Gamma: density per crevice length (per cm, occupied crevices) | | | | | | |
| Large concrete | — | Small concrete | 0.76 | 0.13 | -1.63 | 0.1030 |
| Large concrete | — | Small clay | 0.58 | 0.09 | -3.55 | **0.0012** |
| Small concrete | — | Small clay | 0.77 | 0.11 | -1.83 | 0.0999 |

Table S7: Pairwise contrasts for crevice depth by module type (occupied crevices only) from the Gamma interaction generalized linear mixed model. Recruitment was standardized by crevice area (recruits per cm²) and by crevice length (recruits per cm). Ratio = response ratio (first/second depth). p-values BH-adjusted within each module type. Bold p-values indicate statistical significance.

| Module type | Contrast | Ratio | SE | z | p (BH) |
|---|---|---|---|---|---|
| Density per cm² (occupied crevices) | | | | | |
| Large concrete | Crevice depth 2 — Crevice depth 3 | 2.204 | 0.6182 | 2.82 | **0.0145** |
| Large concrete | Crevice depth 2 — Crevice depth 4 | 1.384 | 0.3701 | 1.22 | 0.2240 |
| Large concrete | Crevice depth 3 — Crevice depth 4 | 0.628 | 0.1761 | -1.66 | 0.1460 |
| Small concrete | Crevice depth 1 — Crevice depth 2 | 1.281 | 0.4331 | 0.73 | 0.8590 |
| Small concrete | Crevice depth 1 — Crevice depth 3 | 1.449 | 0.4902 | 1.1 | 0.8590 |
| Small concrete | Crevice depth 1 — Crevice depth 4 | 1.425 | 0.4819 | 1.05 | 0.8590 |
| Small concrete | Crevice depth 2 — Crevice depth 3 | 1.132 | 0.3315 | 0.42 | 0.8590 |
| Small concrete | Crevice depth 2 — Crevice depth 4 | 1.113 | 0.3259 | 0.36 | 0.8590 |
| Small concrete | Crevice depth 3 — Crevice depth 4 | 0.983 | 0.2879 | -0.06 | 0.9530 |
| Small clay | Crevice depth 1 — Crevice depth 2 | 1.513 | 0.3404 | 1.84 | 0.1320 |
| Small clay | Crevice depth 1 — Crevice depth 3 | 1.691 | 0.3715 | 2.39 | 0.1010 |
| Small clay | Crevice depth 1 — Crevice depth 4 | 1.086 | 0.2444 | 0.37 | 0.7140 |
| Small clay | Crevice depth 2 — Crevice depth 3 | 1.118 | 0.2379 | 0.52 | 0.7140 |
| Small clay | Crevice depth 2 — Crevice depth 4 | 0.718 | 0.1567 | -1.52 | 0.1930 |
| Small clay | Crevice depth 3 — Crevice depth 4 | 0.642 | 0.1366 | -2.08 | 0.1120 |

| | | | | | |
|---|---|---|---|---|---|
| Density per cm (occupied crevices) | | | | | |
| Large concrete | Crevice depth 2 — Crevice depth 3 | 1.676 | 0.4762 | 1.82 | 0.1040 |
| Large concrete | Crevice depth 2 — Crevice depth 4 | 0.895 | 0.2423 | -0.41 | 0.6810 |
| Large concrete | Crevice depth 3 — Crevice depth 4 | 0.534 | 0.1516 | -2.21 | 0.0813 |
| Small concrete | Crevice depth 1 — Crevice depth 2 | 0.905 | 0.3102 | -0.29 | 0.7710 |
| Small concrete | Crevice depth 1 — Crevice depth 3 | 0.742 | 0.2544 | -0.87 | 0.5770 |
| Small concrete | Crevice depth 1 — Crevice depth 4 | 0.557 | 0.1909 | -1.71 | 0.3050 |
| Small concrete | Crevice depth 2 — Crevice depth 3 | 0.82 | 0.2434 | -0.67 | 0.6050 |
| Small concrete | Crevice depth 2 — Crevice depth 4 | 0.615 | 0.1826 | -1.64 | 0.3050 |
| Small concrete | Crevice depth 3 — Crevice depth 4 | 0.75 | 0.2227 | -0.97 | 0.5770 |
| Small clay | Crevice depth 1 — Crevice depth 2 | 1.069 | 0.2438 | 0.29 | 0.7690 |
| Small clay | Crevice depth 1 — Crevice depth 3 | 0.866 | 0.1928 | -0.65 | 0.6220 |
| Small clay | Crevice depth 1 — Crevice depth 4 | 0.425 | 0.0968 | -3.76 | **0.0005** |
| Small clay | Crevice depth 2 — Crevice depth 3 | 0.81 | 0.1746 | -0.98 | 0.4920 |
| Small clay | Crevice depth 2 — Crevice depth 4 | 0.397 | 0.0878 | -4.18 | **0.0002** |
| Small clay | Crevice depth 3 — Crevice depth 4 | 0.49 | 0.1057 | -3.31 | **0.0019** |

Table S8: Pairwise contrasts from the negative binomial generalized linear model comparing planar recruit density (per $m^2$) across module types at Patch Reef 13. Ratio = density ratio (first/second group). p-values adjusted using the Benjamini-Hochberg procedure. Bold p-values indicate statistical significance.

| Contrast | | | Ratio | SE | z | p (BH) |
|---|---|---|---|---|---|---|
| Large concrete | — | Small concrete | 0.89 | 0.26 | -0.49 | 0.6260 |
| Large concrete | — | Small clay | 0.44 | 0.40 | -4.60 | <0.0001 |
| Small concrete | — | Small clay | 0.50 | 0.11 | -3.23 | 0.0019 |

Table S9: Members of the R3D Consortium

| Author | Institution | ORCID |
| --- | --- | --- |
| Benjamin A. Jones | Applied Research Laboratory at the University of Hawaiʻi | 0009-0000-2692-7443 |
| Joshua Levy | Applied Research Laboratory at the University of Hawaiʻi | |
| Sean Mahaffey | Applied Research Laboratory at the University of Hawaiʻi | |
| Aricia Argyris | Applied Research Laboratory at the University of Hawaiʻi | |
| Mark Aruda | Applied Research Laboratory at the University of Hawaiʻi | |
| Ian Robertson | Applied Research Laboratory at the University of Hawaiʻi | |
| Zhenhua Huang | University of Hawaii at Manoa | 0000-0001-6665-7230 |
| Ayrton Medina-Rodriguez | University of Hawaii at Manoa | 0000-0002-0666-9472 |
| Mert Gokdepe | University of Hawaii at Manoa | |
| Brady Halvorson | University of Hawaii at Manoa | |
| Jon Chase | University of Hawaii at Manoa | |
| Charlotte White | University of Hawaii at Manoa | |
| Cami Dillon | University of Hawaii at Manoa | |
| Kristian McDonald | University of Hawaii at Manoa | |
| Anna Mikkelsen | University of Hawaii at Manoa | |
| Josh Madin | Hawaiʻi Institute of Marine Biology | 0000-0002-5005-6227 |
| Mollie Asbury | Hawaiʻi Institute of Marine Biology | |
| Jessica Reichert | Hawaiʻi Institute of Marine Biology | 0000-0003-2245-4188 |
| Hendrikje Jorissen | Hawaiʻi Institute of Marine Biology | |
| Rob Toonen | Hawaiʻi Institute of Marine Biology | 0000-0001-6339-4340 |
| Christopher R. Suchocki | Hawaiʻi Institute of Marine Biology | 0000-0001-6811-1987 |
| Van Wishingrad | Hawaiʻi Institute of Marine Biology | 0000-0002-6256-4018 |
| Chris Jury | Hawaiʻi Institute of Marine Biology | |
| Daniel Schar | Hawaiʻi Institute of Marine Biology | |
| Madeleine Hardt | Hawaiʻi Institute of Marine Biology | |
| Claire Lewis | Hawaiʻi Institute of Marine Biology | 0000-0003-1081-2734 |
| Claire Bardin | Hawaiʻi Institute of Marine Biology | |
| Joshua Kualani | Hawaiʻi Institute of Marine Biology | |
| Crawford Drury | Hawaiʻi Institute of Marine Biology | 0000-0001-8853-416X |
| Kira Hughes | Hawaiʻi Institute of Marine Biology | |
| Josh Hancock | Hawaiʻi Institute of Marine Biology | 0000-0002-4814-1385 |

| | | |
|---|---|---|
| Carlo Caruso | Hawai‘i Institute of Marine Biology | |
| Andrea Grottoli | Ohio State University | 0000-0001-6053-9452 |
| Shannon Dixon | Ohio State University | 0009-0007-9882-7936 |
| Ann Marie Hulver | Ohio State University | 0000-0003-0466-9070 |
| Joshua D. Voss | Florida Atlantic University, Harbor Branch Oceanographic Institute | 0000-0002-0653-2767 |
| Allison Klein | Florida Atlantic University, Harbor Branch Oceanographic Institute | 0009-0004-2670-4222 |
| Siddhartha Verma | Florida Atlantic University, Department of Ocean and Mechanical Engineering | 0000-0002-8941-0633 |
| Alejandro Alvaro | Florida Atlantic University, Department of Ocean and Mechanical Engineering | |
| Richard Argall | Makai Ocean Engineering | |
| Kevin Chun | Makai Ocean Engineering | |
| William Hicks | Makai Ocean Engineering | |
| Alex LeBon | Makai Ocean Engineering | |
| John Yeh | Makai Ocean Engineering | |
| Aaron Thode | Scripps Institution of Oceanography, UC San Diego | |
| Oceane Boulais | Scripps Institution of Oceanography, UC San Diego | |
| Daniel Wangpraseurt | Scripps Institution of Oceanography, UC San Diego | |
| Samapti Kundu | Scripps Institution of Oceanography, UC San Diego | |
| Natalie Levy | Scripps Institution of Oceanography, UC San Diego | |
| Lindsey Badder | Scripps Institution of Oceanography, UC San Diego | |
| Stefan Kolle | Scripps Institution of Oceanography, UC San Diego | |
